# Structural compromise in spiking cortex and connected networks

**DOI:** 10.1101/2024.10.19.619207

**Authors:** Ella Sahlas, Tamir Avigdor, Alexander Ngo, Jessica Royer, Sara Larivière, Judy Chen, Ke Xie, Yezhou Wang, Jack Lam, Raúl Rodríguez-Cruces, Thaera Arafat, Raluca Pana, Andrea Bernasconi, Neda Bernasconi, Birgit Frauscher, Boris C. Bernhardt

## Abstract

**INTRODUCTION:** Epilepsy is increasingly conceptualized as a network disorder, and advancing methods for its diagnosis and treatment requires characterizing both the epileptic generator and related networks. We combined multimodal magnetic resonance imaging (MRI) and high-density electroencephalography (HD-EEG) to interrogate alterations in cortical microstructure, morphology, and intrinsic local function within and beyond spiking tissue in focal epilepsy.

**METHODS:** We studied 25 patients with focal epilepsy (12F, mean ± SD age = 31.28 ± 9.30 years) and 55 age- and sex-matched healthy controls, subdivided into a group of 30 for imaging feature normalization (15F, 31.40 ± 8.74 years) and a group of 25 for replication (12F, 31.04 ± 5.65 years). The 3T MRI acquisition included T1-weighted, diffusion, quantitative T1 relaxometry, and resting-state functional imaging. Open-access MRI processing tools derived cortex-wide maps of morphology and microstructure (cortical thickness, mean diffusivity, and quantitative T1 relaxometry) and intrinsic local function and connectivity (timescales, connectivity distance, and node strength) for all participants. Multivariate approaches generated structural and functional alteration scores for each cortical location. Using HD-EEG electrical source imaging, the most prominent spike type was localized and we quantified MRI alterations within spike sources, as well as in proximal and connected networks.

**RESULTS:** Regions harboring spike sources showed increased structural MRI alterations compared to the rest of the brain in patients. Structural compromise extended to all regions with close functional coupling to spike sources, but not to anatomical neighbors of spike sources. This finding was replicated using average control functional and anatomical matrices instead of patient-specific matrices.

**CONCLUSION:** Spiking regions contain more marked alterations in microstructure and morphology than the remaining cortex, and combining imaging with neurophysiology techniques may ultimately help identify the epileptogenic zone non-invasively. There are nevertheless broader networks effects, which may relate to a cascading of structural changes to functionally connected cortices. These results underscore the utility of combining high-definition MRI and EEG approaches for characterizing epileptogenic tissue and assessing distributed network effects.

## INTRODUCTION

Epilepsy is increasingly conceptualized as both a structural and functional disorder, and advancing its diagnosis and treatment requires characterizing epileptic lesions as well as related networks^1,2^. This is mirrored in the methods used to study epilepsy, of which high-field MRI and HD-EEG are among the most advanced, providing unprecedented spatial and temporal precision^3,4^. Surprisingly, these two modalities have rarely been combined despite their potential synergy in identifying epileptic tissue. Translation of recent advances in non-invasive neuroimaging into clinically viable methods promises to facilitate epilepsy diagnosis, support surgical planning, and elucidate focal epilepsies deemed non-lesional using conventional tools^1,2,5,6,7^. Localizing lesions and networks that harbor epileptogenic activity is especially critical to deliver adequate care to patients with pharmaco-resistant seizures, which affect over 30% of people with epilepsy^8,9^. Poor seizure control reduces quality of life and increases the risk of premature death^8,10,11^. In pharmaco-resistant epilepsy, surgical resection of the region responsible for seizures—the epileptogenic zone—is the most effective treatment to control seizures and improve quality of life^12,13,14^, yet it remains critically underutilized. Only about 1% of potential surgical candidates undergo epilepsy surgery each year in high-income countries^15,16^, and even fewer are operated in middle- and low-income countries^17^. While pharmaco-resistant epilepsy syndromes can often be successfully treated via targeted surgical resections, these conditions also involve large-scale brain networks, with potential implications for surgical prognosis^5^.

A major factor explaining the underutilization of epilepsy surgery lies in the challenge of safely and precisely localizing the epileptogenic zone^2^. This is notably due to the paucity of quantitative markers for the epileptogenic zone that are non-invasive, accurate, and effective. Improved characterization of the epileptogenic zone would be useful clinically, especially in cases considered “imaging-negative”. Neurophysiological work has indicated that localization of interictal epileptic spikes via HD-EEG electrical source imaging can localize the epileptogenic zone^4^. Interictal spikes are more frequent, stable, and safe markers than seizures, and HD-EEG offers more complete brain coverage than intracranial recordings, with no complication risks^18^. Meanwhile, functional MRI studies indicate widespread changes in both local and network function. Recent work in temporal lobe epilepsy suggests marked decreases in intrinsic neural timescales bilaterally^19^. Long-range connectivity, measured with connectivity distance metrics, has also been found to be reduced, mainly ipsilateral to the seizure focus^20^. Node strength, a measure of global network integration, has been shown to co-localize with epileptogenicity in certain patients^21^. Beyond these functional effects, there are increasingly detailed accounts of structural alterations in gray and white matter in focal epilepsy. These involve changes in cortical morphology and microstructure, notably bilateral cortical thickness changes extending beyond the affected lobe in temporal and extra-temporal epilepsy^22,23^, together with changes in quantitative MRI proxies of cortical myelination in temporal lobe epilepsy^24,25^. These findings are complemented by data showing perturbations in the integrity of cortico-subcortical diffusivity, indicating microstructural alterations, in several epilepsy syndromes^26,27,28^.

Despite a rich literature describing electrical, hemodynamic, and structural alterations in epilepsy, there is a lack of studies systematically profiling the neurophysiological epileptic focus and connected regions. Current characterization of epileptic focus and associated networks, thus, remains subject to several limitations. First, many studies assessing alterations across the entire brain in epilepsy do not test whether alterations are most prominent in patient-specific, EEG-defined epileptic foci^19,20,22,24,26^. Additional research is needed to determine whether epileptic foci located via EEG are epicenters of alterations in microstructure, morphology, and intrinsic local function. Second, previous research has not systematically examined brain regions with functional, structural, and anatomical relationships to EEG-defined epileptic foci^4,21^. Consistent with the recognition of epilepsy as a network disorder, the magnitude of alterations across brain regions may relate to their degree of coupling to epileptic generators. Third, much of the existing literature does not capitalize on combinations of modalities to optimize both spatial and temporal resolution in mapping epileptogenicity. Prior MRI research has structurally profiled disease epicentres defined via imaging^28,29^ rather than electrical activity. Here, we address these gaps by quantifying microstructural, morphological, and functional changes measured via high-field MRI within and beyond epileptic spike sources defined by HD-EEG. We further quantify epilepsy-related alterations in adjacent regions as well as functionally and structurally connected neighbors of spike sources to investigate network-level effects.

## METHODS

### Participants

We recruited 25 adult patients diagnosed with focal epilepsy who underwent both multimodal 3T MRI and HD-EEG at the Montreal Neurological Institute and Hospital. The patient cohort included 12 women and 13 men, with a mean age ± standard deviation (SD) of 31.28 ± 9.30 years. The diagnoses at discharge from the epilepsy monitoring unit indicated that the source of seizures was temporal in 14 cases, frontal in 5, fronto-temporal in 3, and temporo-occipital in 3. Twelve cases were right-hemispheric and 13 cases were left-hemispheric. Thirteen patients later underwent epilepsy surgery. Nine patients had an Engel IA outcome at follow-up, two had a IB outcome, one had a IIIA outcome, and one had a IVB outcome. In all operated patients, there was lobe and laterality concordance between the spike source and the resection (8 patients) or thermocoagulation (5 patients). We also recruited 55 age- and sex-matched healthy adults as control participants, divided into two independent control cohorts. The control cohort whose data served for normalization of MRI features (*n* = 30) included 15 women and 15 men, with a mean age ± SD of 31.40 ± 8.74 years. The control cohort whose data served for replication of findings (*n* = 25) included 12 women and 13 men, with a mean age ± SD of 31.04 ± 5.65 years. Control participants underwent the same multimodal 3T MRI as patients and had no history of neurological or psychiatric conditions.

### MRI data acquisition

MRI data were obtained in patients and control participants using a 3T Siemens Magnetom Prisma Fit scanner with a 64-channel head coil receiver. We acquired T1-weighted (T1w) scans via a three-dimensional magnetization-prepared rapid gradient-echo (3D-MPRAGE) sequence (repetition time (TR) = 23000 ms, echo time (TE) = 3.14 ms, inversion time (TI) = 900 ms, flip angle = 9º, matrix = 320 × 320, voxel size = 0.8 × 0.8 × 0.8 mm^3^, field of view (FOV) = 256 × 256 mm^2^, 224 sagittal slices). Diffusion-weighted imaging (DWI) data were obtained using a two-dimensional spin-echo-based echo-planar imaging sequence with multi-band acceleration, based on three shells with b-values of 300 (10 diffusion weighting directions), 700 (40 diffusion directions), and 2000 (90 diffusion directions) s/mm^2^ (TR = 3500 ms, TE = 64.40 ms, flip angle = 90º, refocusing flip angle = 180º, voxel size = 1.6 × 1.6 × 1.6 mm^3^, FOV = 224 × 224 mm^2^, slice thickness = 1.6 mm, multi-band factor = 3, echo spacing = 0.76 ms). Quantitative T1 relaxometry was acquired via a 3D-MP2RAGE sequence (TR = 5000 ms, TE = 2.90 ms, T1 1 = 940 ms, T1 2 = 2830 ms, flip angle 1 = 4º, flip angle 2 = 5º, matrix = 320 × 320, voxel size = 0.8 × 0.8 × 0.8 mm^3^, FOV = 256 × 256 mm^2^, 240 sagittal slices, echo spacing = 7.2 ms, bandwidth = 270 Hx/pixel). Resting-state functional MRI (rsfMRI) was acquired using a two-dimensional blood-oxygen-level-dependent (2D-BOLD) echo-planar sequence with multi-band acceleration (TR = 600 ms, TE = 30 ms, flip angle = 52º, matrix = 80 × 80, voxel size = 3 × 3 × 3 mm^3^, FOV = 240 × 240 mm^2^, slice thickness = 3 mm, multi-band factor = 6, echo spacing = 0.54 ms, bandwidth = 2084 Hz/pixel, 48 slices oriented to the bicommissural line minus 30 degrees). For the ∼7-minute duration of the rsfMRI scan, participants fixated a cross displayed in the centre of the screen and were instructed to not fall asleep.

### MRI data processing

3T MRI data were processed using Micapipe (version 0.2.0, http://micapipe.readthedocs.io)^30^ to extract surface-based measures of cortical microstructure, morphology, and local function (**Figure 1A**).

**Figure 1.**
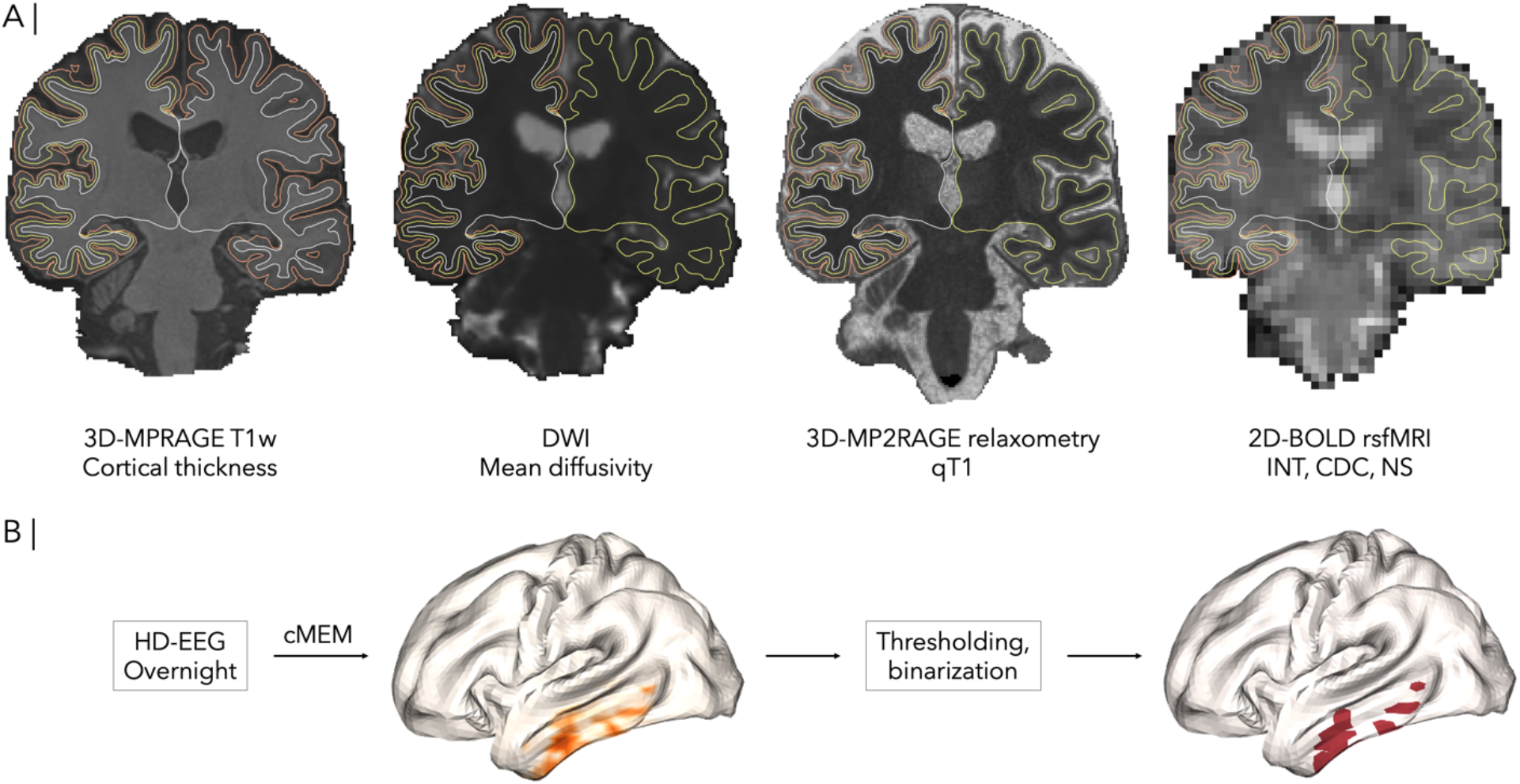
Overview of procedures for multimodal feature extraction. (**a**) Synopsis of multiparametric 3T MRI data acquisition, processing, and features of interest. Acquisition of 3D-MPRAGE, DWI, 3D-MP2RAGE, and 2D-BOLD sequences. Processing using open access software tools: Micapipe (version 0.2.0) and FreeSurfer (version 6.0). Extraction of quantitative features: cortical thickness, mean diffusivity, T1 relaxation time; rsfMRI metrics of intrinsic neural timescale, connectivity distance coefficient, and node strength. Surface-based sampling of cortical thickness between the pial and white matter surfaces; mean diffusivity, quantitative T1 relaxation time, and rsfMRI metrics at the mid-thickness surface. Surfaces are overlaid on the volumetric images of the same patient as in Figure 2. (**b**) Synopsis of HD-EEG data acquisition and processing. Overnight HD-EEG recording using an 83-electrode system. Signal processing using cMEM, then thresholding and binarization to extract vertices with signal amplitude > 50% of the mean at time of maximum spike amplitude for localization of spike sources.

T1w scans were reoriented to standard orientation, deobliqued, corrected for intensity nonuniformity, intensity normalized, and skull stripped. We generated cortical surface models from native T1w scans for each patient using FreeSurfer (version 6.0, https://surfer.nmr.mgh.harvard.edu)^31^. Manual quality control was conducted to correct for segmentation errors in surface extraction via placement of control points and manual edits. Vertex-wise native maps of cortical thickness (CT) were generated in native surface space for each patient by computing the Euclidian distance separating corresponding pial and white matter vertices.

Using MRtrix, DWI images were denoised, b0 intensity-normalized, and corrected for susceptibility distortion, head motion, and eddy currents^32^. Native maps of diffusion tensor-derived mean diffusivity (MD) were generated via interpolation of MD along the surface.

For qT1 relaxometry, 14 equivolumetric surfaces were generated between the pial matter and the white matter boundaries, and qT1 image intensities were sampled systematically using these surfaces. We, thus, obtained vertex-wise profiles of quantitative T1 relaxation time (qT1).

Using Connectome Workbench^33^, we registered native cortical features (CT, MD, and qT1) to the fsLR-5k template surface, with 4842 vertices per hemisphere. We applied spatial smoothing (Gaussian kernel, full width at half maximum (FWHM) = 10 mm) to the resulting CT, MD, and qT1 surface maps.

We processed rsfMRI via AFNI (https://afni.nimh.nih.gov) and FSL (https://fsl.fmrib.ox.ac.uk/fsl), including image reorientation, correction for motion and distortion, removal of nuisance variable signal, averaging of volumetric timeseries, boundary-based registration to native FreeSurfer space, and mapping to surface space using trilinear interpolation^34,35^. Spatial smoothing was also applied (Gaussian kernel, FWHM = 10 mm). To ensure magnetic field saturation, we discarded the first five volumes in each rsfMRI scan.

For each patient, we derived a vertex-wise functional connectivity (FC) matrix by cross-correlating rsfMRI time series, a vertex-wise structural connectivity (SC) matrix from DWI using MRtrix, and a vertex-wise geodesic distance (GD) matrix from T1w imaging using Dijkstra’s algorithm, as previously described^30^. We applied Fischer’s r-to-z transformation to the correlation coefficients in the FC matrix.

Vertex-wise maps of intrinsic neural timescale (INT) were computed for each patient from rsfMRI timeseries using an autocorrelation function^19^. Maps of connectivity distance coefficient (CDC) were generated by thresholding the FC to retain the top 10% of connections, then correlating GD coefficients with the thresholded FC coefficients^20^. Node strength (NS) maps were computed by summing the weights of all connections for each vertex in the FC^21^.

### HD-EEG data acquisition and processing

HD-EEG recordings were acquired overnight during the patients’ hospitalization for pre-surgical evaluation in the epilepsy monitoring unit. Recordings were acquired using a Nihon Koden system (Tokyo, Japan) with 83 electrodes glued with collodion, placed according to the 10–10 EEG system, and sampled at 1000 Hz^36^. Impendences were kept below 1 kΩ for the selected electrodes. The signal was filtered using a Butterworth band-pass filter between 0.3 Hz and 70 Hz. Electrode positions were digitized on the scalp of the patients using a Polhemus localizer device (Vermont, USA). Then, the electrodes were aligned on the head model using reference points (nasion, right, and left ear). Co-registration was refined via a surface fitting approach using Brainstorm^37^. The lead field matrix was computed using the boundary element method with 3 layers for brain, skull, and scalp (with conductivities of 0.33, 0.0165, and 0.33 S/m)^36^, using OpenMEEG implemented in Brainstorm.

### Spike source localization

An experienced epileptologist (B. F.) marked spikes at their peak. The average predominant spike type was localized at the time of maximum amplitude using coherent maximum entropy on the mean (MEM)^38^. MEM is a nonlinear distributed technique for solving the EEG inverse problem. The reference model for coherent MEM is built using a data-driven clustering of the cortical surface into K parcels^4^. We used previously derived FreeSurfer segmentations. To estimate coefficients quantifying the contribution of each dipolar source to the data, we employed the multivariate source pre-localization method^39^. This allowed modelling of the probability of each parcel being active. The ability of coherent MEM to capture the spatial extent of underlying generators has been thoroughly assessed in prior work^4^. Nearest-neighbor interpolation propagated signal probability maps to the same fsLR-5k template surface as the multiparametric MRI data. In each patient, the spike source was defined as vertices with a signal amplitude > 50% of the mean amplitude at the time of maximum spike amplitude, resulting in a binarized map (**Figure 1B**).

### Mapping MRI alteration scores

In each patient, we computed vertex-wise Mahalanobis distances comparing the multivariate aggregate of CT, MD, and qT1 values against corresponding values in 30 control participants. We thus obtained a vertex-wise structural alteration score (**Figure 2A**). Likewise, we computed a functional alteration score based on the Mahalanobis distance aggregating INT, CDC, and NS in patients relative to controls (**Figure 2B**).

**Figure 2.**
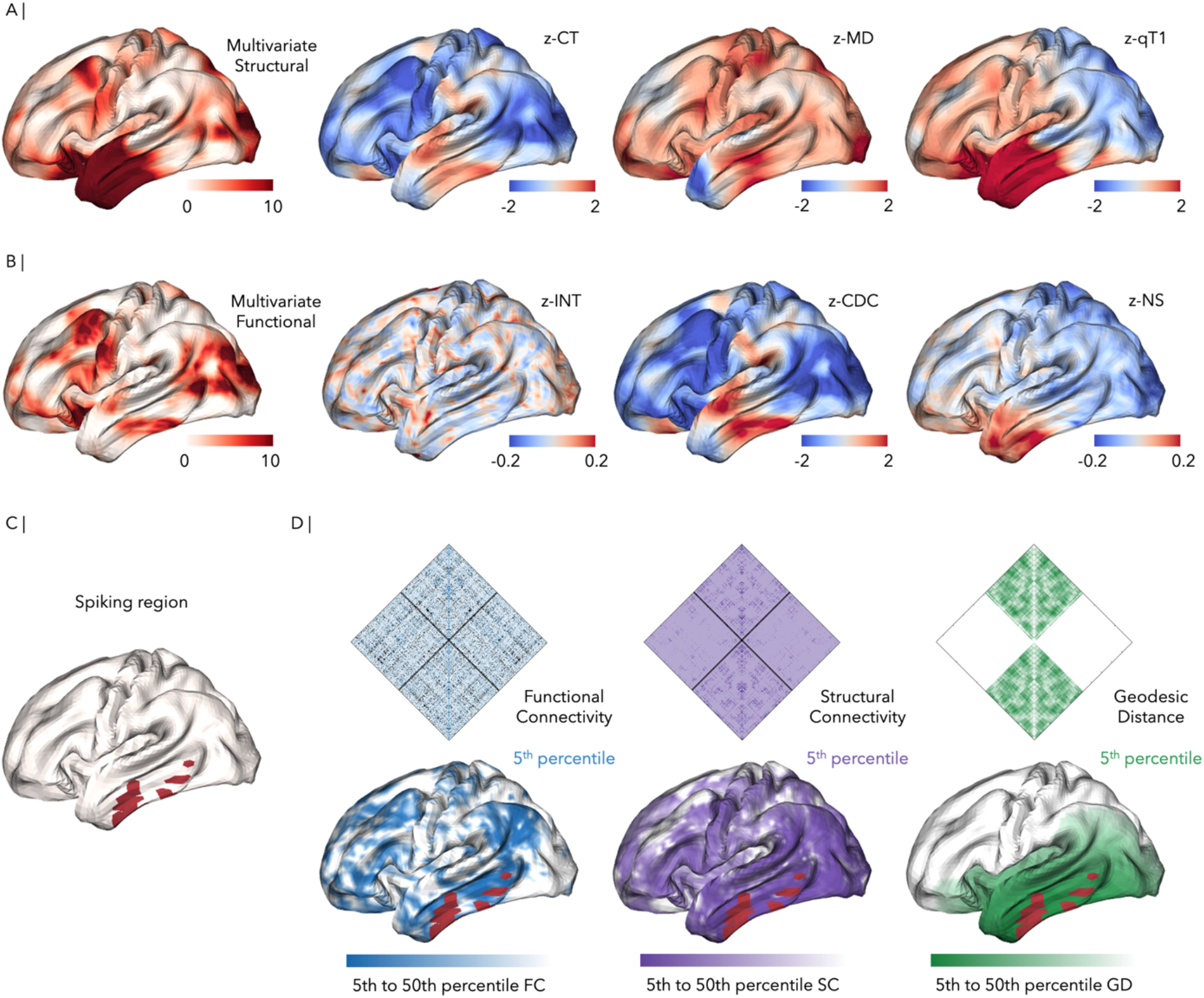
Multivariate scoring of structural and functional alterations and identification of functional, structural, and anatomical neighbors of spike sources. (**a**) Patient-specific map of the multivariate score for alterations in cortical microstructure and morphology, computed as the Mahalanobis distance of cortical thickness (CT), mean diffusivity (MD), and quantitative T1 relaxation time (qT1) at each vertex in the patient relative to the control cohort. For illustration: univariate map of vertex-wise alterations in CT, computed as the z-score of CT at each vertex relative to the control cohort; univariate map of vertex-wise alterations in MD and univariate map of vertex-wise alterations in qT1, computed similarly. Each map is shown in the same representative patient. (**b**) Patient-specific map of the multivariate score for alterations in local functional properties of the cortex, computed as the Mahalanobis distance of intrinsic neural timescale (INT), connectivity distance coefficient (CDC), and node strength (NS) at each vertex relative to the control cohort. For illustration: univariate map of vertex-wise alterations in INT, computed as the z-score of INT at each vertex relative to the control cohort; univariate map of vertex-wise alterations in CDC and univariate map of vertex-wise alterations in NS, computed similarly. Each map is shown in the same representative patient. (**c**) Spike source localized using HD-EEG via coherent maximum entropy on the mean in the same patient. (**d**) Vertices with the 5^th^ to 50^th^ percentiles of functional coupling weights to the spike source (*left*), vertices with the 5^th^ to 50^th^ percentiles of structural connectivity strength to the spike source (*middle*), and vertices with the 5^th^ to 50^th^ percentiles of anatomical proximity to the spike source (*right*).

### Mapping spike sources and their functional, structural, and anatomical neighbors

From the FC, SC, and GD matrices of each patient, we extracted the functional coupling, structural connectivity, and geodesic distance profiles of the spike source relative to all vertices in the rest of the cortex. For each patient, we identified all vertices with above-median functional coupling, structural connectivity, and geodesic proximity to the spike source. We grouped these vertices into ten groups with weights within the 5^th^ to 50^th^ percentile of functional coupling, structural connectivity, and geodesic proximity to the spike source (**Figure 2D**). For assessment of replication, we repeated this procedure using the averaged FC, SC, and GD matrices of the independent cohort of 25 control participants.

### Quantifying MRI alterations in HD-EEG spike sources and their neighbors

We compared the percentage of vertices with significantly altered structural and functional Mahalanobis scores within the spike source to that of the rest of the cortex. The percentage of structurally altered vertices within spike sources was computed as the percent of outlier Mahalanobis distances for CT, MD, and qT1 among all vertices within the spike source. Outlier Mahalanobis distances were defined at the 5% level. Similarly, the percentage of structurally altered vertices in the rest of the cortex was calculated as the percent of outlier CD, MD, and qT1 Mahalanobis distances among all vertices outside of the spike source. The percentages of functionally altered vertices within spike sources and in the rest of the brain was computed as in the structural analysis, using percents of outlier Mahalanobis distances for INT, CDC, and NS. Additionally, we compared structural alteration multivariate scores within regions highly functionally coupled, structurally connected, or geodesically close to the spike source to scores across the rest of the cortex, using patient-specific FC, SC, and GD matrices. We replicated these comparisons using the normative matrices. Comparisons were performed using paired-samples *t*-tests. False Discovery Rate (FDR) corrections were applied to control for multiple comparisons.

### Standard protocol approvals, registrations, and patient consents

#### Research ethics board approval

The Research Ethics Board of the Montreal Neurological Institute and Hospital approved this research involving human participants.

#### Informed consent

All participants in the study provided written informed consent.

### Data availability

Preprocessed cortical thickness, diffusion, and quantitative relaxometry maps used in the study will be made available on the Open Science Framework upon publication.

## RESULTS

### Structural MRI alteration analysis within and beyond HD-EEG spike sources

Cortex within patient-specific spike sources was compared to cortex beyond these sources across all 25 patients (**Figure 3A**). This comparison revealed a greater percentage of structurally altered vertices within epileptic spike sources than across the rest of the cortex (mean: 27.98% versus mean: 17.67%; paired *t*-test; *t* = 2.28; *p* = 0.036; *d* = 0.446).

**Figure 3.**
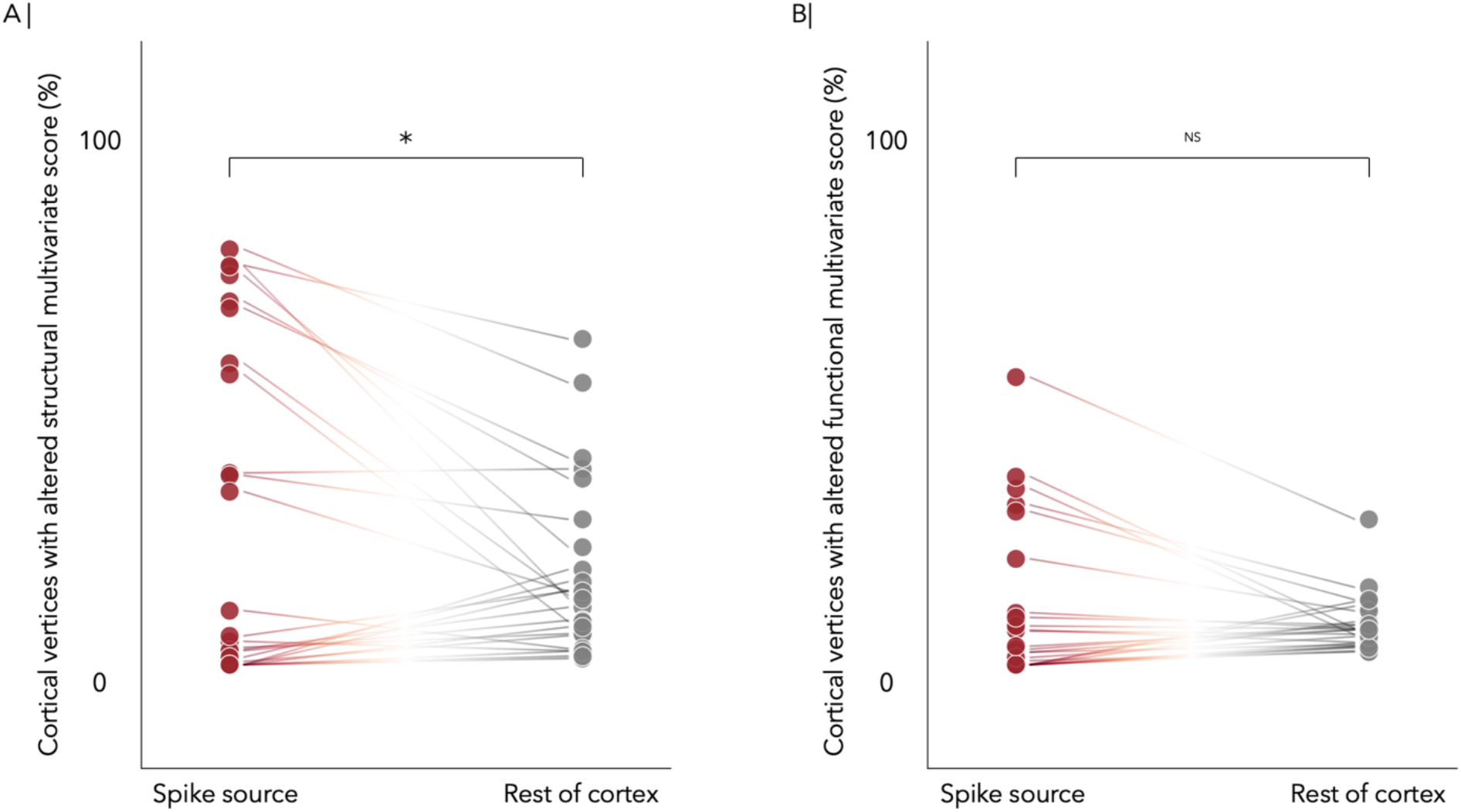
Structural and functional profiling of cortex within spike sources. (**a**) Comparison of the percentage of structurally altered vertices within patient-specific spike sources and in cortex beyond these sources. (**b**) Comparison of the percentage of functionally altered vertices within patient-specific spike sources and in cortex beyond these sources.

We assessed whether electro-clinical focus location was related to structural alterations within spike sources. Thirteen patients had a greater percentage of structural alterations within the spike source. These patients presented with heterogeneous lobar distribution of electro-clinical foci, with foci in temporal (7), frontal (4), and fronto-temporal (2) regions. In 7 of these patients, foci were left-hemispheric, and in 6, foci were right-hemispheric. The 12 patients who did not have a greater percentage of alterations within the spike source had electro-clinical foci in temporal (7), frontal (1), fronto-temporal (1), temporo-occipital (2), and occipital (1) regions; half were left-lateralized and half were right-lateralized. Concordance between structural alterations and HD-EEG spike sources was thus independent of lateralization (χ^2^ test; χ^2^ = 0.00; *p* = 1.00) and lobar distribution of the electroclinical focus (χ^2^ test; χ^2^ = 5.10; *p* = 0.165).

### Functional MRI alteration analysis within and beyond HD-EEG spike sources

There was a subtle increase in the percentage of functionally altered vertices in spike sources compared to the rest of the cortex, but the percentages did not differ significantly (mean: 10.37% versus mean: 7.67%; paired *t*-test; *t* = 1.15; *p* = 0.263; *d* = 0.229; **Figure 3B**).

### Structural alteration analysis in functional, structural, and anatomical neighbors of spike sources

To determine whether the structural compromise identified within spike sources may extend to connected regions, we compared cortex within functional, structural, and anatomical neighbors of spike sources to the rest of the cortex. We located all vertices beyond the spike source that had above-median functional, structural, or anatomical proximity to the spike source. For each of the three neighbor types, we then partitioned all of these vertices into ten groups with decreasing proximity to the spike source. We quantified alterations within each of these ten graded groups of vertices for every neighbor type.

Among the ten groups of vertices with above-median functional coupling to the spike source, all showed higher percentages of structurally altered vertices than the rest of the cortex (paired *t*-tests; *t* ranged from 2.35-3.75; *p*_*FDR*_ ranged from 0.006-0.028). The mean percentages of structurally altered vertices in regions with highest to lowest functional coupling to the spike source are depicted in **Figure 4A** (*left*).

**Figure 4.**
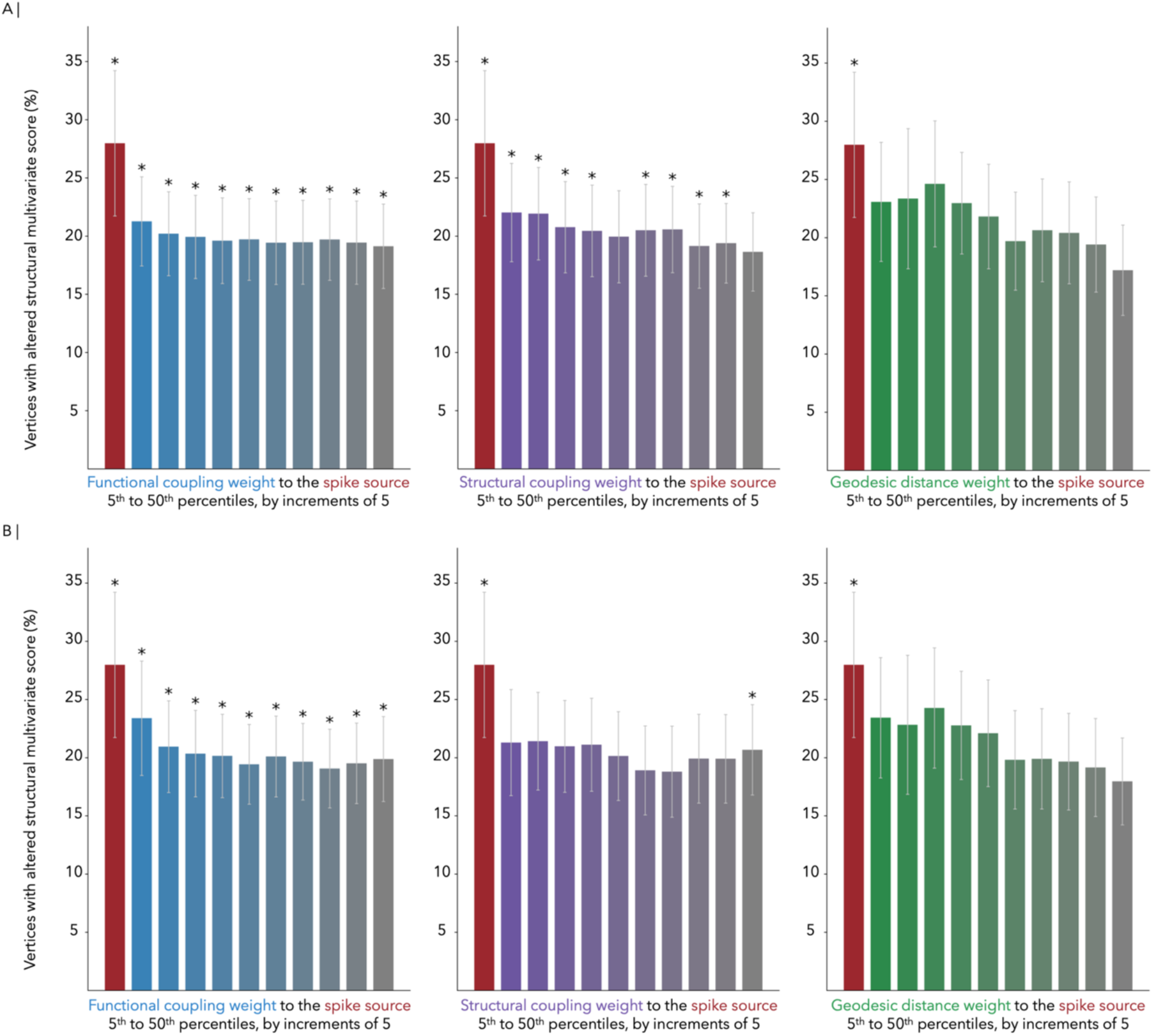
Structural alterations in neighbors of spike sources. (**a**) Quantification of the percentage of structurally altered vertices within functional coupling neighbors (*left*), structural connectivity neighbors (*middle*), and anatomical neighbors (*right*) of spike sources, and comparison to cortex beyond spike sources. Among the ten groups of vertices with above-median functional coupling to the spike source, all showed significantly higher percentages of structurally altered vertices than the rest of the cortex, paired *t*-tests; *t* ranged from 2.35-3.75; FDR-corrected *p* ranged from 0.006-0.028. The mean percentages of structurally altered vertices ranged from 19.13-21.27%. Of the ten groups of vertices with above-median structural connectivity to the spike source, eight showed significantly higher percentages of structurally altered vertices than the rest of the cortex and two were trending, paired *t*-tests; *t* = 1.97-4.36; FDR-corrected *p* = 0.002-0.060. The mean percentages of structurally altered vertices were 18.65-22.03%. Among the ten groups of vertices with above-median anatomical proximity to the spike source, none showed significantly higher percentages of structurally altered vertices than the rest of the cortex, paired *t*-tests; FDR-corrected *p* > 0.05. (**b**) Replication via healthy control (normative) functional coupling, structural connectivity, and geodesic distance: quantification of the percentage of structurally altered vertices within normative functional coupling neighbors (*left*), normative structural connectivity neighbors (*middle*), and normative anatomical neighbors (*right*) of spike sources, and comparison to cortex beyond spike sources. Using normative functional connectivity, all ten groups of vertices with above-median functional coupling to the spike source showed significantly higher percentages of structurally altered vertices than the rest of the cortex, paired *t*-tests; *t* = 2.25-3.82; FDR-corrected *p* = 0.008-0.034. The mean percentages of structurally altered vertices were 19.07-23.39%. With normative structural connectivity, one of the ten groups of vertices with above-median structural connectivity to the spike source contained a significantly higher percentage of structurally altered vertices than the rest of the cortex, paired *t*-tests; *t* = 0.83-3.29; FDR-corrected *p* = 0.031-0.413. The mean percentages of structurally altered vertices were 18.80-21.42%. Of the ten groups of vertices with above-median normative anatomical proximity to the spike source, none showed significantly higher percentages of structurally altered vertices than the rest of the cortex, paired *t*-tests; FDR-corrected *p* > 0.05.

Of the ten groups of vertices with above-median structural connectivity to the spike source, eight showed higher percentages of structurally altered vertices than the rest of the cortex (paired *t*-tests; *t* ranged from 1.97-4.36; *p*_*FDR*_ ranged from 0.002-0.060). **Figure 4A** (*middle*) details the mean percentages of structurally altered vertices in regions with the most to the fewest structural connections to the spike source.

Of the ten groups of vertices with above-median anatomical proximity to the spike source, none showed higher percentages of structurally altered vertices than the rest of the cortex (paired *t*-tests; *t* ranged from

-0.28-2.32; *p*_*FDR*_ ranged from 0.199-0.778), as illustrated in **Figure 4A** (*right*).

### Replication analysis substituting patient-specific connectivity for normative connectivity matrices

Lastly, we assessed whether structural alteration findings in neighbors in spike sources could be replicated using average healthy control functional coupling, structural connectivity, and anatomical distance matrices instead of patient-specific matrices. Normative matrices were generated from the independent cohort of healthy control participants.

Using normative functional connectivity, all ten groups of vertices with above-median functional coupling to the spike source showed higher percentages of structurally altered vertices than the rest of the cortex (paired *t*-tests; *t* ranged from 2.25-3.82; *p*_*FDR*_ ranged from 0.008-0.034), as in **Figure 4B** (*left*).

With normative structural connectivity, only one of the ten groups of vertices with above-median structural connectivity to the spike source contained a higher percentage of structurally altered vertices than the rest of the cortex (paired *t*-tests; *t* ranged from 0.83-3.29; *p*_*FDR*_ ranged from 0.031-0.413). The mean percentages of structurally altered vertices in these regions are represented in **Figure 4B** (*middle*).

Among the ten groups of vertices with above-median normative anatomical proximity to the spike source, none showed higher percentages of structurally altered vertices than the rest of the cortex (paired *t*-tests; *t* ranged from 0.21-2.33; *p*_*FDR*_ ranged from 0.241-0.837), as shown in **Figure 4B** (*right*).

In sum, structural alteration findings were replicated in functional and anatomical neighbors, but not structural connectivity neighbors, across two methods of computing functional coupling, structural connectivity, and anatomical distance matrices: the patient-specific approach and an approach averaging matrices across healthy control participants. Both methods were consistent with the presence of structural alterations in functional neighbors of spike sources, and the lack thereof in anatomical neighbors.

## DISCUSSION

Capitalizing on multiple imaging features and on both spatial and temporal detail, our findings chart a map of focal epilepsies characterized by *(i)* increased structural—but not functional—compromise captured by MRI within EEG-defined spike sources, and *(ii)* extension of structural compromise to regions functionally coupled to—but not solely anatomically nearby—spike sources. Taken together, these findings incite additional multimodal studies analyzing both epileptogenic tissue and related networks. Results also highlight the potential for synergy between MRI and HD-EEG for identifying epileptogenic tissue, which may help advance non-invasive and seizure-independent epilepsy diagnostics. Unlike previous research using EEG-fMRI^40^, our study focuses on microstructure and thus combines modalities without performing simultaneous EEG and MRI, as the high-quality microstructural MRI acquisitions would be technically challenging and uncomfortable for patients in the context of overnight HD-EEG recordings. Notably, prior investigations have improved the accuracy of non-invasive approaches in localizing epileptogenicity by combining modalities^40,41,42^. Nonetheless, epilepsy research has more commonly remained siloed across neurophysiology and imaging domains. Consequently, while neurophysiology research has developed increasingly reliable methods for localizing epileptogenic activity, whether through iEEG^43^ or HD-EEG^4^, MRI research has focused on mapping lesions, which do not always coincide with the epileptogenic zone^42^. There is a marked paucity of studies investigating structural MRI measures in EEG-defined epileptogenic regions, yet approaches of this kind are requisite for deciphering the structural makeup of cortex harboring seizures. This is illustrated in the use of both structural imaging and stereotactic EEG data in the generation of personalized virtual brain models^2^. Morphological alterations have also been found to modulate circuit excitability in computational models of epilepsy^44^, which may drive circuits to generate epileptic activity. Multimodal approaches combining MRI and HD-EEG also carry potential for clinical impact. HD-EEG offers multiple opportunities for non-invasively localizing the epileptogenic zone^4,41^, while high-field MRI is unparalleled in detecting subtle, epilepsy-causing lesions^45,46,47^. Together, these modalities hold potential to locate the surgical target in cases where conventional methods may not contribute to diagnosis and surgical target definition. This combination could also ultimately help limit or more precisely target invasive pre-surgical investigations in an increasing number of patients with pharmaco-resistant epilepsy.

The present study suggests that regions harboring EEG spike sources contain MRI-derived microstructural and morphological alterations. This is in line with studies that have applied machine learning methods to localizing epileptogenic lesions based on data extracted from MRI^45,46,47^. Much of this literature has focused on automating the detection of focal cortical dysplasia, which often still eludes conventional MRI evaluations^45^. To validate localization approaches, prior studies have trained models such as feed-forward artificial neural networks on morphometric maps^45^ and deep convolutional neural networks on T1w and fluid-attenuated inversion recovery images^46^. While MRI-based localization approaches have become increasingly successful, several outstanding avenues remain to be explored. First, studies have rarely validated MRI approaches against electro-clinical evidence. Instead, studies commonly use lesions segmented from MRI as the ground truth. This introduces potential confounds as methods are developed and tested using the same MRI data. While some studies have validated approaches against surgical cavity information or histopathological data^47^, additional validation based on spike sources defined electrically could provide further insights into intrinsic epileptic activity. The approach presented in our study could provide context for the integration of neurophysiological benchmarks into future studies aiming to validate MRI-based detection algorithms. Second, accuracy rates of automated lesion localization currently average at around 80%, at most^45,46,47^. Careful feature selection and weighting may introduce clinically meaningful improvements in accuracy rates. The present study is motivated by prior work highlighting epilepsy-related changes in cortical thickness, relaxometry, and diffusion parameters^22,24,26^. Our findings are consistent with increased alterations in these features within the epileptic focus, providing grounds for their inclusion in future lesion detection work. Of note, in our study, the spike source was defined at spike peaks to optimize temporal resolution and signal-to-noise ratio. While this approach may identify the time of maximal cortical involvement most precisely, analyzing the spike midpoint can be preferred to prioritize localizing spike onset prior to early propagation^36^. Our study does not find evidence of significantly increased alterations in MRI-sensitive intrinsic function within spike sources. Additional studies, ideally in larger samples, may be required to detect possible changes in MRI-derived metrics of intrinsic function localized to spike sources. Consistent with existing evidence, alterations in intrinsic function may be widely distributed across the cortex^19,20^ and less concentrated within spike sources relative to structural changes, thus, differences across spiking and non-spiking tissue may be more subtle and difficult to detect.

Prior research has characterized network-related disruptions in epilepsy, collectively contributing to the growing recognition of pharmaco-resistant epilepsies as network disorders^19,20,22,24,26,48^. Indeed, alterations in intrinsic neural timescales, cortical thickness, and diffusion properties have been shown to extend bilaterally in patients with temporal lobe epilepsy and in populations suffering from other epilepsy syndromes^19,22,26^. Recent work in temporal lobe epilepsy has identified widespread age-related changes in cortical thickness and diffusion metrics, extending beyond the ipsilateral mesiotemporal lobe^48^. Prior investigations have also identified alterations in connectivity distance and relaxometry extending to ipsilateral regions beyond the seizure focus^20,24^. Still, little work has tested whether alteration epicenters correspond to connectivity neighbors of the epileptic focus. Certain studies have assessed epilepsy-related disruptions in structural networks^49^ and evaluated the impact of connectivity profiles on atrophy patterns^50^. However, no study to date has quantified structural alteration patterns with functional and anatomical relationships to the electrically defined spike source. Our study addresses this gap via a novel method for multiparametric alteration scoring, in cortex with varying connections to the spike source. Our findings are consistent with the absence of increased alterations in the geodesic neighbors of spike sources, indicating that the patterning of the microstructural signature of epileptic activity may not be explained simply by anatomical proximity to spike sources. In our study, alterations were more frequently identified in functional than in structural neighbors of spike sources, suggesting that regions of microstructural damage may be linked by polysynaptic rather than monosynaptic connections. Longitudinal structure-function studies are needed to ascertain whether this reflects pre-existing structural irregularities in functional networks or alterations induced over time by the propagation of chronic seizures. Previously described disease-specific changes in structural networks^49^ may explain the contrasting presence of alterations in most patient-specific structural connectivity neighbors and lack thereof in most normative structural connectivity neighbors. Our study identified increased microstructural alterations in regions with above-median functional coupling to regions harboring spike sources, suggesting a possible cascading of microstructural alterations to cortical regions beyond spike sources through functional network effects. This finding is consistent with focal epilepsy invading large-scale networks, and underscores the utility of quantitative multimodal MRI to identify otherwise undetectable lesions and help guide surgical planning, in particular when used in combination with electrophysiological data and clinical correlates^7,29,47^.

The present study highlights MRI-derived alterations in cortical microstructure and morphology that co-localize with epileptic spike sources, and that cascade into polysynaptic functional networks in which spike sources are embedded. Intrinsic functional alterations, on the other hand, may be distributed more broadly, with less specific co-localization than structural changes. While results are overall consistent with synergy of HD-EEG and high-field MRI findings in identifying epilepsy-related alterations, additional work is recommended to optimize the weighting of modalities and imaging parameters across epilepsy syndromes and individual patients. These findings will advance current conceptions of focal epilepsy and help to uncover clinically relevant network effects that may impact treatment and prognosis.

## ACKNOWLEDGMENTS

E.S. acknowledges funding from the Fonds de Recherche du Québec – Santé (FRQS) Doctoral Training Scholarship. B.F. acknowledges funding from a project grant from the Canadian Institutes of Health Research (CIHR) (PJT-175056). B.C.B. acknowledges funding from CIHR (FDN-154298, PJT-174995), the Natural Sciences and Engineering Research Council of Canada (NSERC) (Discovery-1304413), FRQS, the Brain Canada Foundation, the SickKids Foundation (NI17-039), the Azrieli Center for Autism Research (ACAR), the Helmholtz International BigBrain Analytics and Learning Laboratory (HIBALL), and the Canada Research Chairs Program.

## AUTHOR CONTRIBUTIONS

E.S. and B.C.B. contributed to conception and design of the study; E.S., T.A., A.N., J.R., S.L., J.C., B.F., and B.C.B. contributed to data acquisition; E.S. contributed to processing of MRI data; T.A. and B.F. contributed to processing of EEG data and writing of EEG methods; E.S. and B.C.B. analyzed the data and wrote the manuscript; all authors revised and approved the manuscript.

## TABLE

**Table 1.** Participant demographics.

|  | **Patient cohort** | **Normalization**<br>**control cohort** | **Replication**<br>**control cohort** |
| --- | --- | --- | --- |
| **Number** | 25 | 30 | 25 |
| **Sex (F/M)** | 12/13 | 15/15 | 12/13 |
| **Age at scan (years)** | 31.28 ± 9.30 | 31.40 ± 8.74 | 31.04 ± 5.65 |
| **Lateralization (L/R)** | 13/12 | N/A | N/A |

